# The IκB-Protein BCL-3 Controls MAPK Activity by Promoting TPL-2 Degradation in the Nucleus

**DOI:** 10.1101/373209

**Authors:** Patricia E. Collins, Domenico Somma, David Kerrigan, Felicity Herrington, Karen R. Keeshan, Robert J. Nibbs, Ruaidhrí J. Carmody

**Author notes:** Corresponding author: Ruaidhrí J. Carmody, Institute of Infection, Immunity and Inflammation, College of Medicine, Veterinary and Life Sciences, University of Glasgow, Glasgow, G12 8TA United Kingdom.

## Abstract

The ability of the innate immune system to distinguish between low level microbial presence and invasive pathogens is fundamental for immune homeostasis and immunity. However, the molecular mechanisms underlying threat discrimination by innate immune cells are not clearly defined. Here we describe the integration of the NF-ĸB and MAPK pathways in the nucleus by the IĸB protein BCL-3 and the MAP3K TPL-2. Our data reveals that TPL-2 is a nucleocytoplasmic shuttling protein and demonstrates that the nucleus is the primary site for TPL-2 ubiquitination and proteasomal degradation. BCL-3 promotes TPL-2 degradation through interaction in the nucleus. As a consequence, *Bcl3*^-/-^ macrophages have increased TPL-2 stability and MAPK activity following TLR stimulation. The enhanced stability of TPL-2 in *Bcl3*^-/-^ macrophages lowers the MAPK activation threshold and the level of TLR ligand required to initiate an inflammatory response. This study establishes the nucleus as a key regulatory site for TLR-induced MAPK activity and identifies BCL-3 as a regulator of the cellular decision to initiate inflammation

## Introduction

Toll-like receptor (TLR) activation is essential for the development of an effective immune response to infection or injury. TLR-induced transcriptional responses are primarily dependent on the activation of the NF-ĸB and MAPK pathways^1,2^. Activation of NF-ĸB leads to the transcription of a large number of pro-inflammatory cytokines, chemokines, and adhesion molecules which activate and mobilise an immune response to the initiating insult^3^. TLR-induced activation of the MAPK pathway controls the expression of a number of transcription factors such as *Fos*, *Egr1* and *Elk1* as well as certain cytokines and immunomodulators^2, 4-6^. MAPK activation also plays a critical role in the post-transcriptional control of cytokine expression through the regulation of mRNA stability, transport and translation^2^. Indeed, NF-ĸB activation in the absence of MAPK activation does not lead to inflammation as increases in pro-inflammatory gene transcription are not accompanied by increased translation^7^. Thus, the coordinated control of these pathways is required to ensure that the inflammatory response is appropriate to the initiating stimulus.

Although TLR activation of the NF-ĸB and MAPK pathways both require the IKKβ kinase^8^, each pathway possesses distinct features that determine the consequences of TLR ligation. IKKβ triggered degradation of IĸBα induces the activation of NF-ĸB proportional to the level of stimulus leading to graded transcriptional responses^9,10^. In contrast, the MAPK pathway is ultrasensitive whereby its activation primarily occurs over a narrow range of ligand concentrations^11,12^. This property of the MAPK pathway effectively permits it to switch from “off” to “on” and enables cells to implement “yes/no” decisions on whether to initiate an inflammatory response based on TLR ligand concentration^13,14^. Importantly, at TLR ligand concentrations below the MAPK activation threshold, NF-ĸB activation is un-coupled from inflammation and instead, NF-κB activity appears to mediate macrophage priming ^7,13^.

The serine/threonine kinase TPL-2 (MAP3K8) is essential for MAPK activation by TLR ligands and cytokines including TNFα and IL1β^15-17^. In resting cells, TPL-2 is bound by NF-ĸB p105 which prevents it from interacting with and phosphorylating MEK1^15,17^. Activation of IKKβ phosphorylates p105 bound to TPL-2 triggering p105 ubiquitination and proteasomal degradation, and liberating active TPL-2 which in turn initiates the MAPK cascade by phosphorylating and activating MEK1^18,19^. Active TPL-2 is highly unstable and is rapidly degraded by the proteasome^15,17^. TPL-2 degradation is the primary mechanism for limiting TLR-induced MAPK activation, indeed, in the absence of p105 (*Nfkb1*) TPL-2 protein levels are extremely low and *Nfkb1*^-/-^ macrophage and mice are deficient in TLR-induced MAPK activation.

Here we show that the IĸB protein BCL-3, which is a negative regulator of NF-ĸB transcriptional activity^20^, is also an important negative regulator of TLR-induced MAPK activity. *Bcl3*^-/-^ macrophages have elevated levels of MAPK activation following TLR stimulation leading to increased expression of MAPK-dependent target genes. We establish TPL-2 as a nucleocytoplasmic shuttling protein and identify a N-terminal nuclear export sequence (NES) sufficient to mediate nuclear export. Our data demonstrate that TPL-2 undergoes ubiquitination in the nucleus and that the nuclear localisation is a critical determinant of TPL-2 half-life. BCL-3 binds TPL-2 to promote its proteasomal degradation in the nucleus and thereby regulates MAPK activity. The role of BCL-3 as a negative regulator of TPL-2 is independent of its role as a negative regulator of NF-ĸB. At the cellular level, BCL-3 mediated inhibition of TPL-2 determines the threshold level of TLR ligand required for a pro-inflammatory response in macrophages, and *Bcl3*^-/-^ macrophages produce cytokines at levels of TLR ligand that do not elicit a response in wild type macrophages. As a consequence, *Bcl3*^-/-^ mice have elevated levels of circulating pro-inflammatory cytokines but not homeostatic chemokines in the resting state. Together, these findings identify BCL-3 as a unique factor linking the regulation of the MAPK and NF-kB pathways in the nucleus and identify BCL-3 as a critical factor in the cellular decision to initiate an inflammatory response.

## Results

### BCL-3 negatively regulates MAPK-dependent gene expression

We previously identified the IĸB protein BCL-3 as a negative regulator of TLR-induced transcriptional responses through the stabilisation of NF-ĸB p50 homodimer repressor complexes^20,21^. On further analysis we observed that in addition to inhibiting the expression of NF-ĸB regulated pro-inflammatory genes, BCL-3 also limits the TLR-inducible transcription of a number of immediate/early genes that rely on MAPK activity for expression^22^. Bone marrow derived macrophages (BMDMs) from wild type (WT) and *Bcl3*^-/-^ mice were left untreated or stimulated with LPS and mRNA analysed by probe directed RNA-seq to measure the core TLR-induced transcriptional response. As expected^20^, *Bcl3*^-/-^ macrophages are hyper-responsive to LPS stimulation as measured by the elevated expression of NF-ĸB target genes including *Tnf*, *Il6* and *Il1b* (Figure 1A). Remarkably, *Bcl3*^-/-^ macrophages also showed significantly increased levels of mRNA for *Fos*, *Egr1*, *N4ra1* and *Dusp5* following LPS stimulation when compared to WT controls (Figure 1A). These data were confirmed by QPCR analysis in independent experiments (Figure 1B). Stimulation of macrophages with other TLR ligands (CpG, PAM3CSK4 and Poly[I:C]) and TNFα also led to increased expression of MAPK target genes in *Bcl3*^-/-^ cells over WT cells (Supplemental Figure S1). Since the TLR-inducible expression of *Fos*, *Egr1*, *N4ra1* and *Dusp5* is strongly dependent on MAPK activity^22^ we further investigated the role of MAPK activity in the enhanced expression of these genes in *Bcl3^-/-^ cells*. WT and *Bcl3*^-/-^ macrophages were stimulated with LPS in the presence of increasing concentrations of the MEK1 inhibitor U0126 and the expression of *Egr1* measured by QPCR. This analysis revealed that the LPS-inducible expression of *Egr1* was MAPK dependent in both WT and *Bcl3*^-/-^ cells and that WT and *Bcl3*^-/-^ macrophages are equally sensitive to MEK1 inhibition (Figure 1C). Similar results were found for LPS-induced expression of *Dusp1*, *Fos*, *Egr2* while the expression of *Tnf* was found to be insensitive to MEK1 inhibition in both WT and *Bcl3*^-/-^ cells (Figure 1D). Together these data identify a new role for BCL-3 in limiting the MAPK-dependent expression of immediate/early genes following TLR stimulation and suggest an NF-ĸB-independent role for BCL-3 in regulating TLR responses.

**Figure 1.**
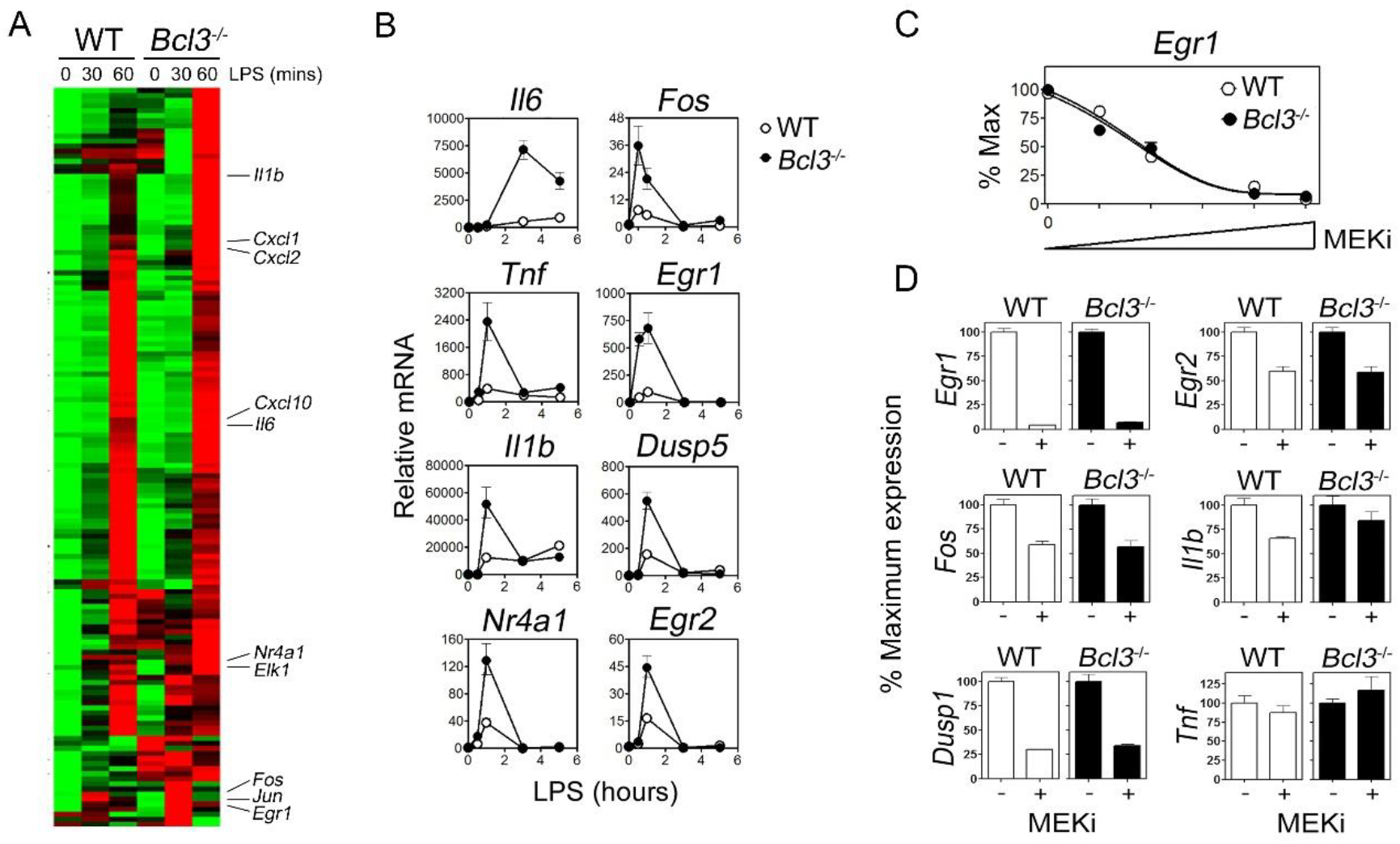
Increased MAPK dependent gene expression in *Bcl3*^-/-^ macrophages. **(A)** Heat map of RNA-seq analysis of wild type (WT) and *Bcl3*^-/-^ macrophages stimulated with LPS (10ng/ml) for the indicated times. **(B)** QPCR analysis of selected genes in WT and *Bcl3*^-/-^ macrophages stimulated with LPS (10ng/ml) for the indicated times. **(C)** QPCR analysis of LPS-induced Egr1 expression (10ng/ml for 60 mins) in WT and *Bcl3*^-/-^ macrophages in the presence of increasing amounts of the MEK1 inhibitor U1026 (MEKi). Relative levels of *Egr1* mRNA are expression as a % of max for each genotype. (**D**) QPCR analysis of selected genes in WT and *Bcl3*^-/-^ macrophages stimulated with LPS (10ng/ml 60mins) in the presence or absence of MEKi (0.5μM). Data representative of 3 independent experiments.

### BCL-3 inhibits TPL-2 activation of the MAPK pathway

The enhanced expression of MAPK-dependent genes in TLR- and TNFR-stimulated *Bcl3*^-/-^ macrophages suggested a defect in the MAPK pathway in the absence of BCL-3. To investigate this we next analysed the activation of the MAPK pathway in LPS stimulated WT and *Bcl3*^-/-^ macrophages using phospho-specific antibodies against MEK1, ERK1/2 and p90RSK. This revealed a significant increase in the strength and duration of the LPS-induced MAPK cascade in *Bcl3*^-/-^ cells compared to WT cells characterised by increased phosphorylation of MEK1, ERK1/2 and RSK (Figure 2A, B and C). The increased LPS-induced activation of ERK1/2 in *Bcl3*^-/-^ cells was sensitive to treatment with MEK1 inhibitor demonstrating that the enhanced activation of the MAPK pathway in *Bcl3*^-/-^ cells results from enhanced MEK1 activity (Figure 2D). TPL-2 (MAP3K8) is the apical kinase of the MAPK pathway activated by TLRs and TNFR and directly phosphorylates and activates MEK1. To investigate the effect of BCL-3 on TPL-2-induced MAPK activity independently of TLR activation we measured the impact of BCL-3 expression on TPL-2 induced AP-1 reporter activity using RAW 264.7 macrophages. Co-expression of BCL-3 blocked TPL-2-induced AP-1 reporter activity while overexpression of the NF-ĸB p50 subunit did not (Figure 2E). Together these data identify BCL-3 as an inhibitor of TLR-induced MAPK activation by TPL-2 kinase. The inhibition of TPL-2 activity by BCL-3 but not NF-ĸB p50 identifies this as a NF-ĸB-independent role of BCL-3, since the negative regulation of transcriptional activity by Bcl-3 is dependent on p50^20,21^.

**Figure 2.**
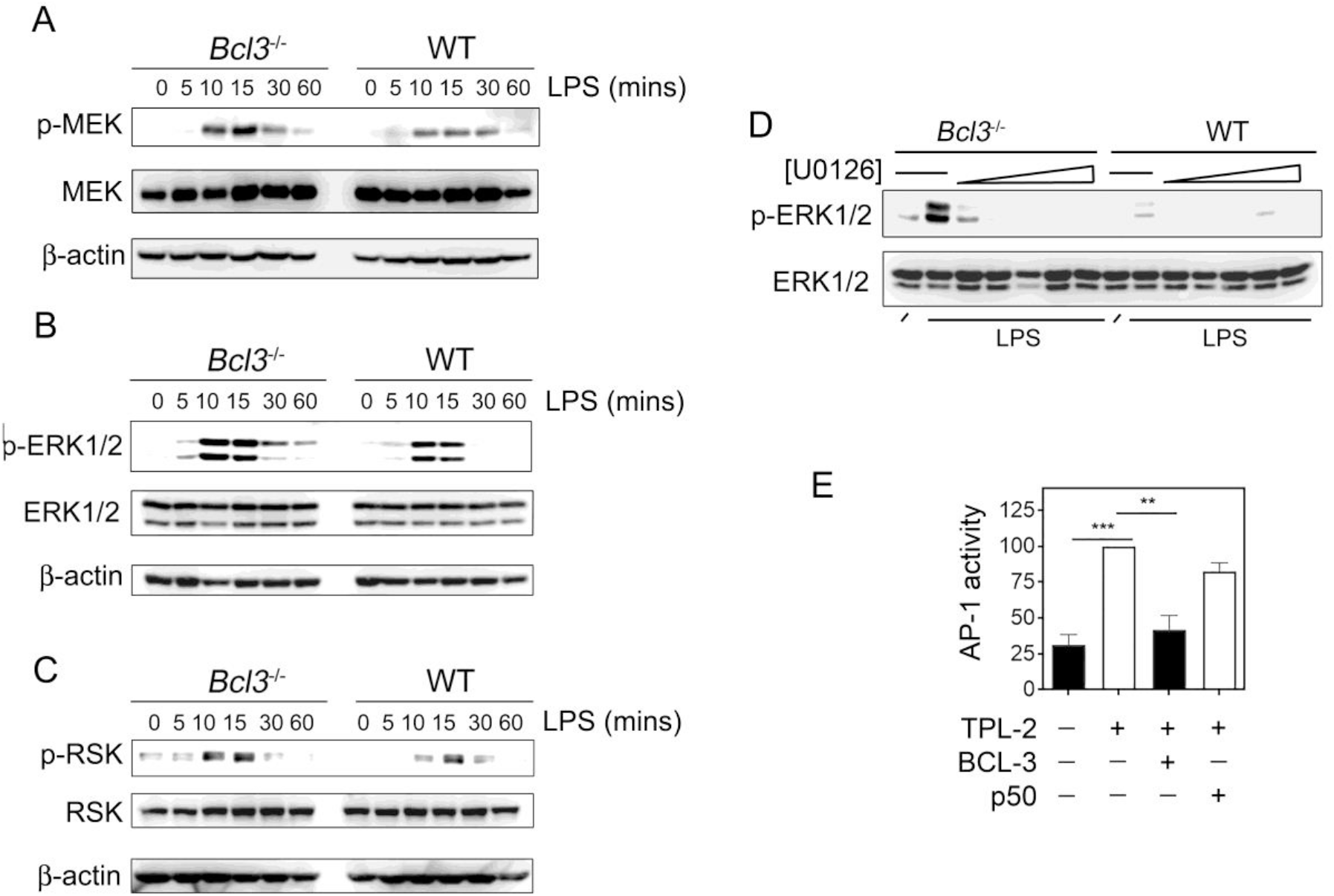
BCL-3 inhibits MAPK activity. WT and *Bcl3*^-/-^ macrophages were stimulated with LPS (10ng/ml) for the indicated times and phosphorylation of MEK1 **(A)**, ERK1/2 **(B)**, and RSK **(C)** analysed by immunoblot. **(D)** WT and *Bcl3*^-/-^ macrophages were cells were pre-treated for 30 mins with MEK1 inhibitor U0126 prior to stimulation with LPS (10ng/ml for 5 mins) in the absence or presence of increasing amounts of the MEK1 inhibitor U0126 (0 - 0.5 μM) and phosphorylation of ERK1/2 analysed by immunoblot. **(E)** RAW macrophage cells were transfected with an AP-1 luciferase reporter plasmid along with expression vectors for TPL-2, BCL-3 or NF-ĸB p50 as indicated and reporter activity measured. Data representative of 3 independent experiments -/+ SEM. *** p>0.001, ** p>0.01 by one way ANOVA.

### BCL-3 interacts with TPL-2

The TLR-induced activation of the MAPK and NF-ĸB pathways is linked *via* NF-ĸB p105 which acts as an inhibitor of both pathways^23^. p105 inhibits MAPK activation by binding to TPL-2 through its C-terminal ankyrin repeat domain^17^. The p105 C-terminal ankyrin repeat domain bears significant structural and sequence homology with the central ankyrin repeat domain of BCL-3 suggesting the potential interaction of BCL-3 with TPL-2. Co-immunoprecipitation experiments using HEK293T cells transiently transfected with plasmids encoding BCL-3 and TPL-2 revealed a readily detectable interaction between BCL-3 and TPL-2 (Figure 3A). Further co-immunoprecipitation experiments demonstrated the interaction of endogenous BCL-3 and TPL-2 in THP-1 monocytes (Figure 3B). NF-ĸB p105 and p50 are not required for the interaction of BCL-3 and TPL-2 since BCL-3 and TPL-2 also co-immunoprecipitated when expressed in *Nfkb1*^-/-^ murine embryonic fibroblast (MEF) cells further supporting the NF-ĸB-independent role of BCL-3 in the regulation of MAPK (Figure 3C). Additional experiments using a kinase inactive mutant of TPL-2 revealed that TPL-2 kinase activity is needed for interaction with BCL-3, a requirement similar to that reported for the interaction of TPL-2 with p105^24^ (Figure 3D). These data raise the possibility that the interaction of BCL-3 with TPL-2 inhibits TPL-2 kinase activity to limit the activation of the MAPK pathway and the subsequent transcriptional response to TLR activation. However, an *in vitro* kinase assay employing purified recombinant BCL-3 and TPL-2 proteins and using recombinant MEK1 as substrate revealed that BCL-3 does not directly inhibit TPL-2 phosphorylation of MEK1 (Figure 3E). Thus BCL-3 is not an inhibitor of TPL-2 kinase activity.

**Figure 3.**
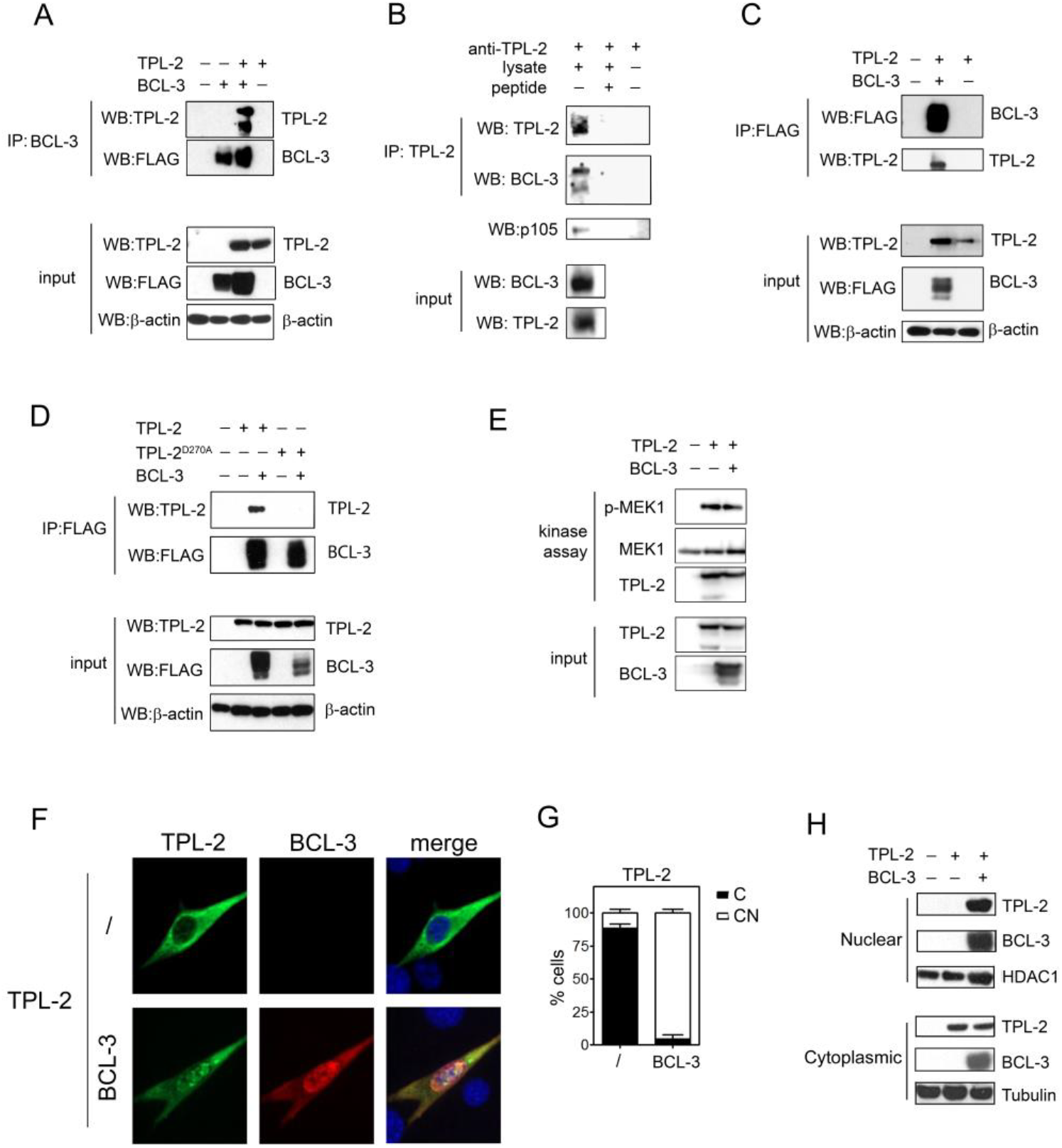
BCL-3 interacts with TPL-2. **(A)** HEK293T cells were transfected with plasmids encoding BCL-3 and TPL-2 prior to immunoprecipitation of BCL-3 and immunoblot using anti-TPL-2 antibody. **(B)** TPL-2 was immunoprecipitated form THP-1 lysates and immunoblotted using anti-BCL-3 antibody. Inclusion of immunising TPL-2 peptide demonstrated specificity of immunoprecipitation. **(C)** *Nfkb1*^-/-^ MEFs were transfected with plasmids encoding BCL-3 and TPL-2. Cells were pre-treated with 20uM MG132 for 2 hours prior to analysis. Immunoprecipitated BCL-3 was immunoblotted with anti-TPL-2 antibody. **(D)** HEK293T cells were transfected with plasmids encoding BCL-3, TPL-2 and a kinase dead mutant of TPL-2 (D270A) prior to immunoprecipitation of BCL-3 and immunoblot using anti-TPL-2 antibody.**(E)** TPL-2 invitro kinase assay in the presence and absence of recombinant BCL-3 using MEK1 as substrate. Phosphorylation of MEK1 was detected by anti-p-MEK1 immunoblot. **(F)** Confocal immunofluorescence microscopy of 3T3 cells transfected with plasmids encoding TPL-2 and BCL-3. DAPI nuclear stain shown in blue. **(G)** Quantification of TPL-2 subcellular localisation. The subcellular distribution of TPL-2 was scored as nuclear and cytoplasmic (NC) or predominantly cytoplasmic (C) and presented as the percentage of total cells counted. Data shown are the mean +/- SEM of 3 independent experiments (n=50 per experiment). **(H)** Nuclear and cytoplasmic fractions of HEK293T cells transfected with plasmids encoding TPL-2 and BCL-3 were analysed by immunoblot using indicated antibodies.

Since TPL-2 is a cytoplasmic protein and BCL-3 is localised in the nucleus we next determined the subcellular localisation of TPL-2 and BCL-3 interaction. Unexpectedly, immunofluorescence microscopy revealed that the interaction of TPL-2 and BCL-3 occurs in the nucleus and that the expression of BCL-3 promotes the subcellular re-distribution of TPL-2 from the cytoplasm to the nucleus (Figure 3F and 3G). BCL-3 induced re-localisation of TPL-2 to the nucleus was further confirmed by immunoblotting of nuclear and cytoplasmic extracts (Figure 3H). These data identify BCL-3 and TPL-2 as novel interaction partners and demonstrate that this interaction takes place in the nucleus.

### TPL-2 is a nucleocytoplasmic shuttling protein

The interaction of BCL-3 with TPL-2 was unexpected as BCL-3 is a nuclear protein while TPL-2 is considered a cytoplasmic protein. Subcellular fractionation demonstrated a small but readily detectable pool of TPL-2 protein in the nuclear fraction of macrophages (Figure 4A). Both cytoplasmic and nuclear levels of TPL-2 protein were rapidly decreased following stimulation of macrophages with LPS, consistent with the rapid proteasomal degradation of active TPL-2. Of note, the LPS-induced degradation of TPL-2 protein occurred with faster kinetics in the nuclear fraction compared to the cytoplasmic fraction suggesting that nuclear TPL-2 is more unstable than cytoplasmic TPL-2.

**Figure 4.**
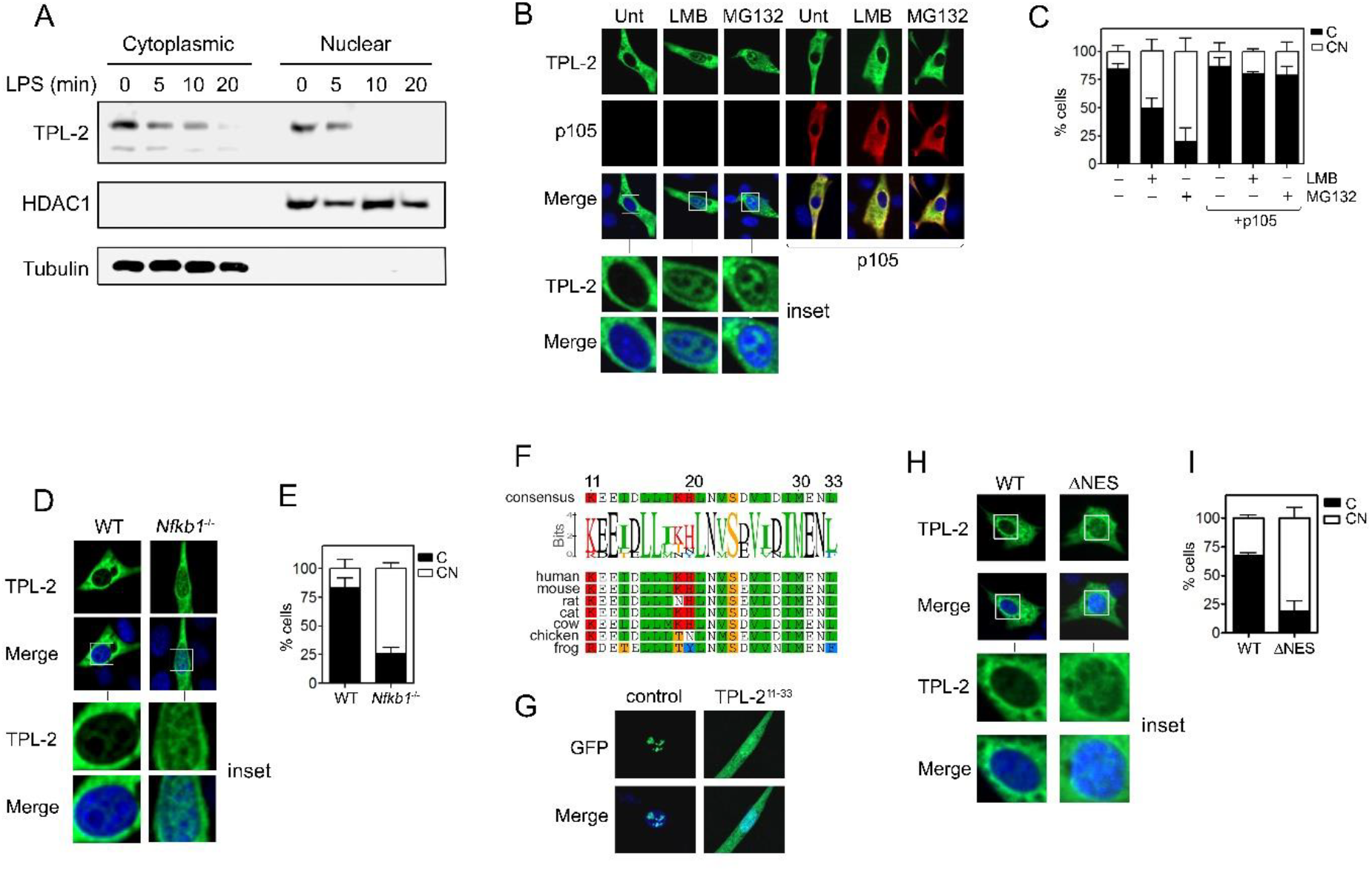
TPL-2 is a nuclear cytoplasmic shuttling protein. **(A)** WT macrophages were stimulated with LPS (10ng/ml) for the indicated times before equal amounts nuclear and cytoplasmic protein fractions were analysed by immunoblot using indicated antibodies. **(B)** Confocal immunofluorescence microscopy of 3T3 cells transfected with plasmids encoding TPL-2 and p105 and treated with leptomycin B (20nM for 2hours) and MG132 (20μM for 2hours). DAPI nuclear stain shown in blue. **(C)** % of cells from (B) displaying cytoplasmic or nuclear and cytoplasmic distribution of TPL-2. **(D)** Confocal immunofluorescence microscopy of *Nfkb1*^-/-^ MEFs cells transfected with plasmid encoding TPL-2. DAPI nuclear stain shown in blue. **(E)** % of cells from (D) displaying cytoplasmic or nuclear and cytoplasmic distribution of TPL-2. **(F)** Identification of a conserved nuclear export sequence (NES) in the N terminal of TPL-2. **(G)** The TPL-2 NES sequence is sufficient to mediate the nuclear export of a REV1-GFP fusion protein. DAPI nuclear stain shown in blue. **(H)** Immunofluorescence microscopy of 3T3 cells transfected with plasmid encoding TPL-2 or a TPL-2 mutant lacking the NES (ΔNES). DAPI nuclear stain shown in blue. **(I)** % of cells from (H) displaying cytoplasmic or nuclear and cytoplasmic distribution of TPL-2. The subcellular distribution of TPL-2 in (B), (D) and (H) was scored as nuclear and cytoplasmic {NC} or predominantly cytoplasmic {C} and presented as the percentage of total cells counted. Data shown are the mean +/- SEM of 3 independent experiments (n=50 per experiment).

Treatment of cells expressing TPL-2 with leptomycin B, an inhibitor of CrmA/Exportin-1 mediated nuclear export, led to significant accumulation of TPL-2 in the nucleus demonstrating that TPL-2 is a nucleocytoplasmic shuttling protein (Figure 4B and 4C). Importantly, similar results were obtained following treatment of cells with the proteasome inhibitor MG132 (Figure 4B and 4C), demonstrating that TPL-2 is highly unstable in the nucleus, findings consistent with LPS-induced degradation of nuclear TPL-2 in macrophages (Figure 4A). Interestingly, co-expression of p105 prevented TPL-2 nuclear localisation induced by both leptomycin B and proteasome inhibition (Figure 4B and 4C). These data indicate that the maintenance of TPL-2 protein levels by p105 may result from the inhibition of TPL-2 nuclear localisation. This was supported by similar experiments using WT and *Nfkb1*^-/-^ MEFs which demonstrated that in cells lacking p105 TPL-2 constitutively localises to the nucleus without the requirement for inhibition of the proteasome or Crm1 mediated nuclear export (Figure 4D and 4E). These findings also strongly correlate with the instability of TPL-2 protein in *Nfkb1*^-/-^ cells.

Analysis of the TPL-2 amino acid sequence identified a conserved N-terminal putative nuclear localisation sequence (NES) predicted to mediate leptomycin B sensitive CrmA/Exportin-1 mediated nuclear export ^25,26^ (Figure 4F). Insertion of the TPL-2 NES sequence into a NES-deficient Rev-GFP fusion protein^27^ was sufficient to export the Rev-GFP protein from the nucleus to the cytoplasm, demonstrating that this region of TPL-2 is a *bone fide* NES (Figure 4G). Deletion of the NES in TPL-2 led to the accumulation of TPL-2 in the nucleus (Figure 4H and 4I) at levels similar to that induced by leptomycin B treatment (Figure 4B) but does not inhibit interaction with p105 or BCL-3 (Supplemental Figure S2). These data establish TPL-2 as a nuclear-cytoplasmic shuttling protein containing an N-terminal NES.

### TPL-2 undergoes ubiquitination and proteasomal degradation in the nucleus

The accumulation of TPL-2 in the nucleus induced by proteasome inhibition suggests that the nucleus is the subcellular site of TPL-2 degradation. In support of this, treatment of TPL-2 expressing cells with leptomycin B led to a dramatic reduction in TPL-2 protein levels which were restored by proteasome inhibition (Figure 5A). Furthermore, deletion of the NES of TPL-2 (TPL-2^ΔNES^) leads to reduced TPL-2 levels compared to full legth TPL-2 which were also restored by proteasome inhibition, demonstrating that nuclear localisation of TPL-2 leads to its proteasomal degradation (Figure 5B). Similar to proteasome inhibition, expression of p105 stabilised both full length TPL-2 and TPL-2^ΔNES^ protein levels (Figure 5C). p105 expression blocked the nuclear accumulation of TPL-2^ΔNES^ protein demonstrating that the cytoplasmic sequestration of TPL-2 by p105 is a key mechanism regulating its stability (Figure 5D and E). An in-cell ubiquitination assay revealed TPL-2 to be highly polyubiquitinated and demonstrated that p105 is a potent inhibitor of TPL-2 polyubiquitination, consistent with its role as a stabiliser of TPL-2 (Figure 5F). Importantly, the vast majority of polyubiquitinated TPL-2 is detected in the nuclear fraction of cellular extracts (Figure 5G). Together these data identify the nucleus as the site of TPL-2 polyubiquitination and proteasomal degradation and that a key mechanism for stabilising TPL-2 protein is the cytoplasmic sequestration of TPL-2 by p105. These data also indicate that BCL-3 may regulate the MAPK pathway through nuclear sequestration and enhanced proteasomal degradation of TPL-2.

**Figure 5.**
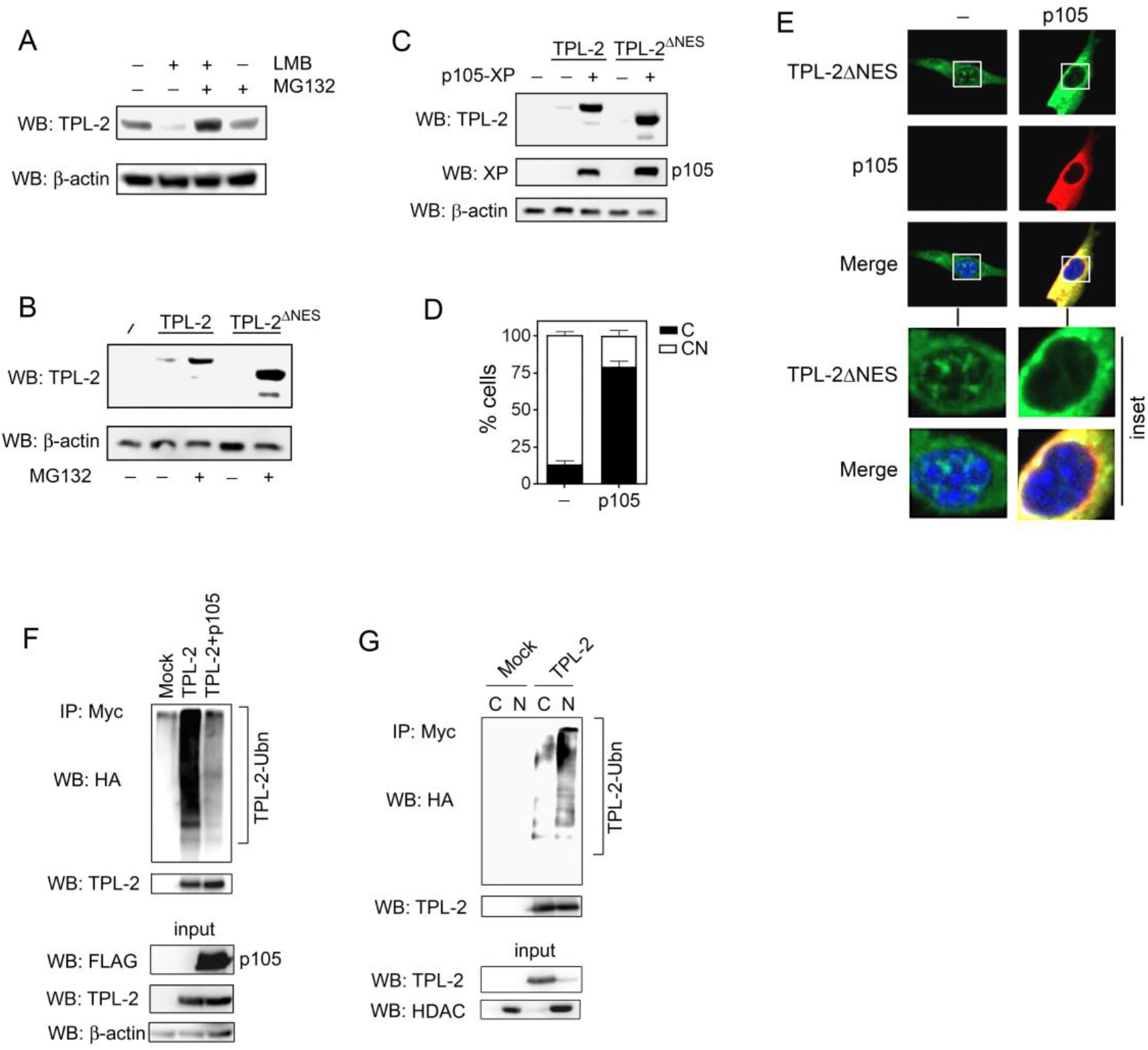
TPL-2 is degraded in the nucleus. **(A)** HEK293T cells transfected with plasmid encoding TPL-2 and treated with LMB (20nM for 6 hours) and MG132 (20μM for 2hours) and analysed by immunoblot using indicated antibodies. **(B)** HEK293T cells transfected with plasmid encoding TPL-2 and TPL^ΔNES^ and treated with MG132 (10μM, 2 hours) before analysis by immunoblot using indicated antibodies. **(C)** HEK293T cells transfected with plasmid encoding TPL-2, TPL^ΔNES^ and p105, and treated with MG132 (10μM, 2 hours) before analysis by immunoblot using indicated antibodies. **(D)** % of cells from (E) displaying cytoplasmic or nuclear and cytoplasmic distribution of TPL-2. Quantification of TPL-2 subcellular localisation. The subcellular distribution of TPL-2 was scored as nuclear and cytoplasmic (NC) or predominantly cytoplasmic (C) and presented as the percentage of total cells counted. Data shown are the mean +/- SEM of 3 independent experiments (n=50 per experiment). **(E)** Immunofluorescence microscopy of 3T3 cells transfected with plasmids encoding TPL-2ΔNES and p105. DAPI nuclear stain shown in blue. **(F)** HEK293T cells were transfected with plasmids encoding Myc-TPL-2, FLAG-p105 and HA-tagged ubiquitin. Lysates were immunoprecipitated with antibody against TPL-2 and immunoblotted with antibody against HA. **(G)** HEK293T cells were transfected with plasmids encoding Myc-TPL-2 and HA-tagged ubiquitin. Nuclear {N} and cytoplasmic {C} fractions were immunoprecipitated with anti TPL-2 antibody and immunoblotted with antibody against HA. Data representative of 3 independent experiments.

### BCL-3 controls TPL-2 nuclear localisation and proteasomal degradation

Our data demonstrates that TPL-2 is a nuclear cytoplasmic shuttling protein which is selectively degraded by the proteasome in the nucleus. Our data also demonstrate that nuclear localisation of TPL-2 is controlled by p105 and BCL-3, suggesting that BCL-3 promotes the degradation of TPL-2 to limit MAPK pathway activity. In support of this, expression of BCL-3 induces TPL-2 degradation which is reversed by proteasome inhibition (Figure 6A). Moreover, the half-life of TPL-2 in LPS stimulated *Bcl3*^-/-^ macrophages is approximately 2 fold greater than that of WT macrophages demonstrating that BCL-3 promotes TPL-2 degradation to regulate MAPK activity (Figure 6B). Importantly, TPL-2 still undergoes LPS induced degradation in *Bcl3*^-/-^ cells, albeit with significantly slower kinetics, demonstrating that BCL-3 is not essential for TPL-2 degradation.

**Figure 6.**
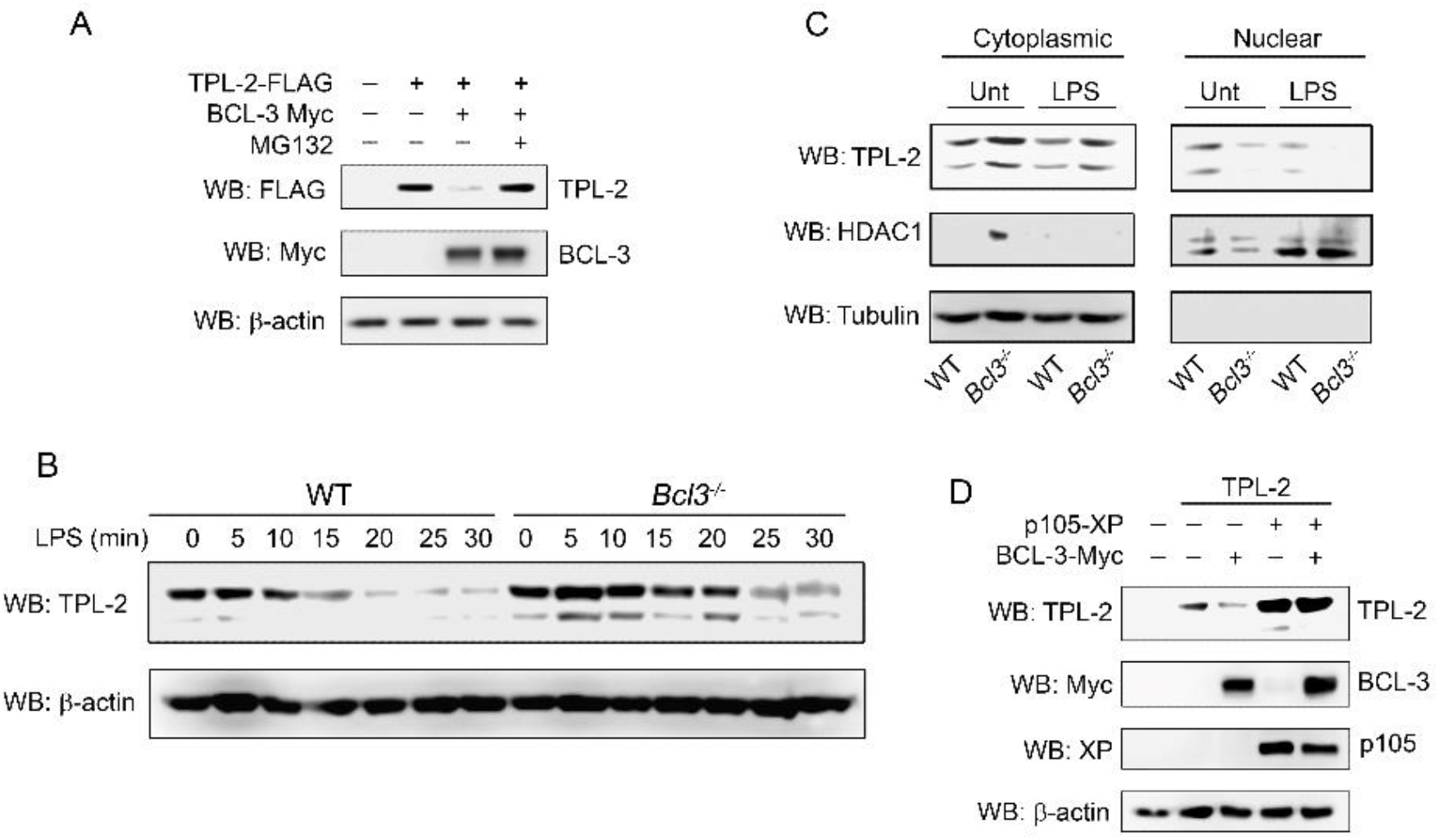
BCL-3 promotes the degradation of TPL-2. **(A)** HEK293T cells transfected with plasmids encoding Myc-BCL-3 and FLAG-TPL-2 and treated with MG132 (10μM for 2 hours). Relative protein levels were determined by immunoblotting using indicated antibodies. **(B)** WT and *Bcl3*^-/-^ macrophages were stimulated with LPS (10ng/ml) for the indicated times prior to analysis by immunoblotting using the indicated antibodies. **(C)** WT and *Bcl3*^-/-^ macrophages were stimulated with LPS (10ng/ml for 10mins) before nuclear and cytoplasmic fractions were analysed by immunoblot using anti-TPL-2 antibody. **(D)** HEK293T cells were transfected with plasmids encoding TPL-2, XP-p105 and Myc-BCL-3. Relative protein levels were determined by immunoblotting.

Low levels of TPL-2 are detectable in the nucleus of WT macrophages which are rapidly lost following LPS stimulation suggesting that TPL-2 degradation in the nucleus may also require signal induced phosphorylation of TPL-2 (Figure 4A). Significantly reduced levels of TPL-2 are found in the nucleus of *Bcl3*^-/-^ macrophages, which instead demonstrate a greater cytoplasmic localisation of TPL-2 in both unstimulated and LPS stimulated conditions (Figure 6C). These data demonstrate that BCL-3 interaction with TPL-2 serves to shift the equilibrium of neucleocytoplasmic shuttling TPL-2 towards the nucleus, thereby increasing the proteasomal degradation of TPL-2 and limiting MAPK activation. In the absence of BCL-3 the altered equilibrium of TPL-2 nuclear cytoplasmic distribution leads to a slower rate of degradation and extended TPL-2 half life. In contrast p105 prevents BCL-3 induced TPL-2 degradation by preventing TPL-2 localisation to the nucleus (Figure 6D).

### BCL-3 sets the activation threshold for LPS-induced responses

The MAPK pathway is ultrasensitive which allows TLRs to effectively switch MAPK activity from “off” to “on” over a narrow range of ligand concentrations to initiate an inflammatory response to stimulation^13^. To determine the role of BCL-3 in regulating the MAPK activation threshold in response to TLR activation we first measured MAPK activity in WT and *Bcl3*^-/-^ macrophages stimulated with a range of LPS concentrations. This demonstrated that *Bcl3*^-/-^ macrophages activate the MAPK pathway at significantly lower concentrations of LPS than WT macrophages (Figure 7A, 7B and S3) suggesting that *Bcl3*^-/-^ macrophages have a lower activation threshold than WT cells. The activation of the NF-ĸB pathway, as measured by IĸBα degradation, was equivalent between WT and *Bcl3*^-/-^ cells (supplemental Figure S4) as was the number of cells responding to stimulation (Supplemental Figure S5). To assess the impact of this altered MAPK activation threshold in *Bcl3*^-/-^ macrophages we next analysed the LPS-induced production of cytokines and chemokines. This revealed production of pro-inflammatory cytokines including TNFα, IL-12 and IL-1β in *Bcl3*^-/-^ macrophages treated with low concentrations of LPS that did not elicit a response in WT cells (Figure 7C and D). Remarkably this effect was selectively associated with pro-inflammatory cytokines while the production of chemokines was equivalent between WT and *Bcl3*^-/-^ macrophages treated with the same concentrations of LPS (Figure 7C and D). Equivalent levels of cytokine mRNA were induced at low concentrations of LPS in both WT and *Bcl3*^-/-^ macrophages indicating that the increased production of cytokines by *Bcl3*^-/-^ macrophages results from post transcriptional regulation of cytokines (Supplemental Figure S6). In support of this we observed increased *Tnf* RNA stability in *Bcl3*^-/-^ macrophages compared to WT cells (Supplemental data Figure S7) a finding consistent with the previously reported role for TPL-2 in the post-transcriptional control of cytokine expression^4-6^. A similar profile of cytokine and chemokine levels was observed in the serum of WT and *Bcl3*^-/-^ mice. *Bcl3*^-/-^ mice showed significantly elevated levels of serum TNFα, IL1β, IL-13 and IL-6 compared to WT mice, while the circulating levels of CXCL1, CCL2, CCL4 G-CSF, and GM-CSF were equivalent between *Bcl3*^-/-^ and WT mice (Figure 7E and F). Administration of LPS *in vivo* however led to significantly elevated serum levels of both cytokines and chemokines in *Bcl3*^-/-^ mice compared to WT mice, consistent with the role of BCL-3 as a negative regulator of inflammatory responses to high levels of LPS (Supplemental Figure S8).

**Figure 7.**
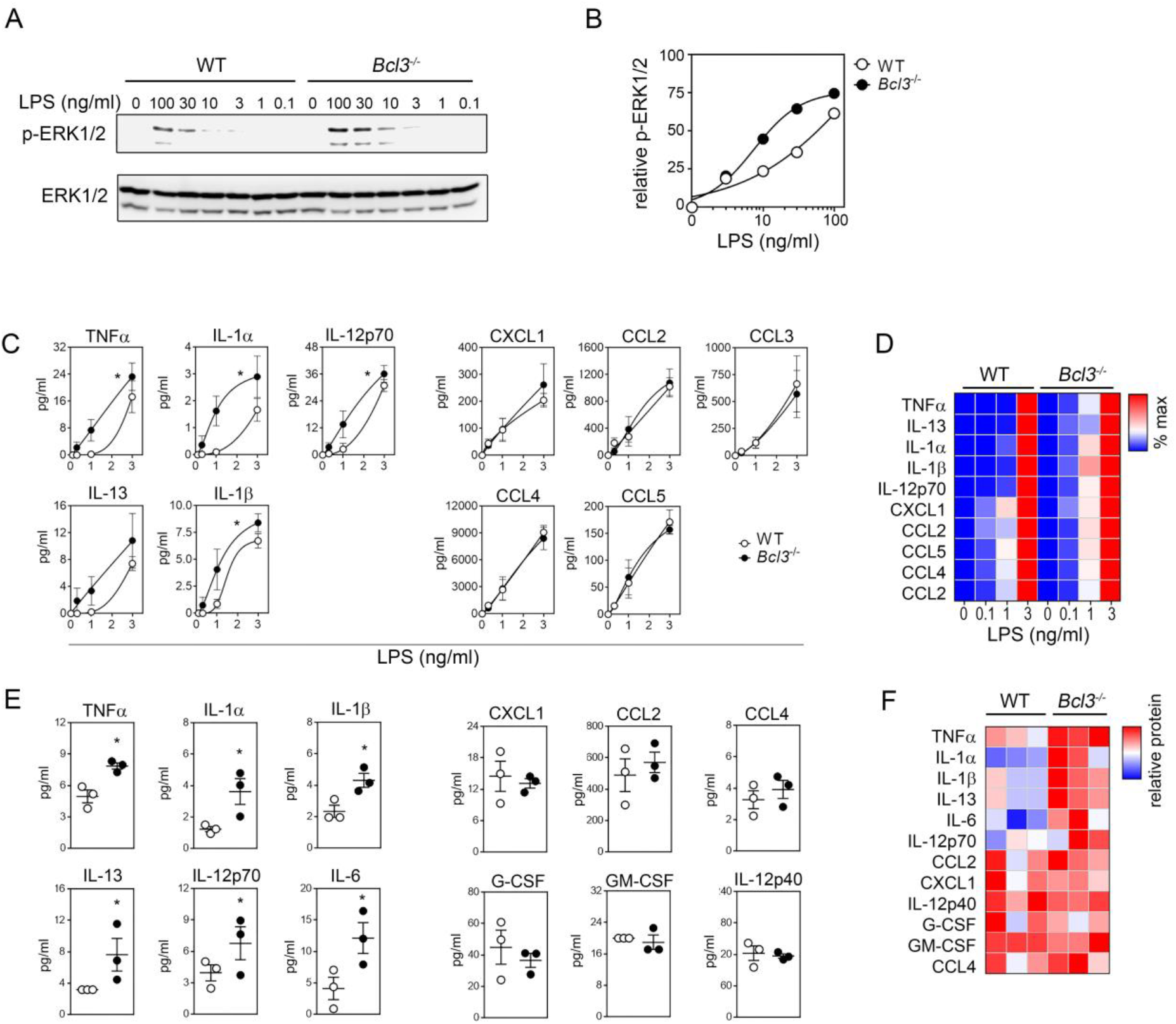
BCL-3 regulates TLR response threshold. **(A)** WT and *Bcl3*^-/-^ macrophages were stimulated with the indicated concentration of LPS and phosphor-ERK1/2 levels measured by immunoblotting. **(B)** Immunoblotting data from (A) was quantitatively analysed and normalised to total ERK1/2 protein levels. **(C)** WT and *Bcl3*^-/-^ macrophages were stimulated with the indicated concentrations of LPS and levels of secreted cytokines and chemokines analysed by Luminex assay. Data is the mean -/+ SEM of 3 independent experiments. **(D)** Relative levels of secreted factors as measured in (C) expressed as a % of maximum for WT and *Bcl3*^-/-^ macrophages. **(E)** Serum from WT and *Bcl3*^-/-^ mice was analysed for the indicated factors by Luminex assay. Data shows the mean -/+ SEM (n=3). **(F)** Relative levels of soluble factors as measured in (E) for WT and *Bcl3*^-/-^ mice.

## Discussion

This study identifies a new role for the IĸB protein BCL-3 in limiting inflammatory responses to TLR activation independently of its previously described role in regulating NF-ĸB transcriptional activity^20^. We establish BCL-3 as a unique factor that integrates the control of the NF-ĸB and MAPK pathways in the nucleus. Our data demonstrates that at low levels of stimulus, BCL-3 primarily functions to inhibit inflammation by negatively regulating MAPK activity while at high levels of stimulus, BCl-3 inhibits NF-ĸB transcriptional activity to limit pro-inflammatory gene expression^20^. Our data shows that BCL-3 not only regulates the level of MAPK activation in response to stimulation, but importantly determines the MAPK activation threshold and the level of TLR ligand required to generate an inflammatory response. The control of MAPK activity by BCL-3 is mediated through TPL-2 kinase, which we reveal is a nucleocytoplasmic shuttling protein that undergoes ubiquitination and proteasomal degradation in the nucleus. BCL-3 interacts with TPL-2 in the nucleus to enhance it proteasomal turnover and limit activation of the MAPK pathway.

Our data establish TPL-2 as a nucleocytoplasmic shuttling protein and identify a conserved C terminal nuclear export sequence in TPL-2 that mediates the nuclear export of TPL-2 through the leptomycin B sensitive Crm1/exportin-1 system. Nuclear TPL-2 is highly unstable and undergoes ubiquitination and proteasomal degradation. Indeed, the nucleus is the primarily site of TPL-2 ubiquitination and degradation. Interaction of TPL-2 with p105 has previously been demonstrated to stabilise TPL-2 by inhibiting TPL-2 proteasomal degradation^15,17^. Our data demonstrates that p105 stabilises TPL-2 by preventing its translocation to the nucleus rather than directly inhibiting TPL-2 ubiquitination and proteasomal degradation. Thus, in unstimulated cells, TPL-2 is stable and cytoplasmic while in activated cells, TPL-2 shuttles between the cytoplasm and the nucleus where it is unstable. The equilibrium of nuclear and cytoplasmic distribution of TPL-2 is determined by the interaction of BCL-3 with TPL-2 which increases the nuclear residence of TPL-2 to increase its turnover and reduce its half-life. BCL-3 is not required for TPL-2 degradation, as TPL-2 is degraded in *Bcl3*^-/-^ macrophages, but rather increases the rate of TPL-2 turnover. Interestingly, TPL-2 is detected in the nucleus of unstimulated macrophages, suggesting that TPL-2 degradation may also be accelerated by TLR activation, potentially by signal -induced phosphorylation.

The proteasomal degradation of TPL-2 controls not only the duration of MAPK activity following TLR activation, but also the MAPK activation threshold – the level of stimulation required to activate MAPK to a sufficient degree to allow the production of pro-inflammatory cytokines^13^. In contrast to the NF-ĸB pathway, in which the activity of NF-ĸB is proportional to the level of TLR stimulation^9,10^, the majority of MAPK activation occurs over a narrow range of TLR ligand concentrations^13^. This characteristic of the MAPK pathway is termed ultrasensitivity, and enables cells to implement “yes” or “no” decisions on whether to initiate an inflammatory response on the basis of TLR ligand concentrations^11^. This study establishes the MAPK activation threshold as a regulatable property, one that may be influenced by factors affecting TPL-2 nuclear localisation and proteasomal degradation. Thus, *Bcl3*^-/-^ macrophages exhibit a much lower activation threshold than wild types cells, leading to significant MAPK activation and cytokine production a very low concentrations of TLR ligand that do not elicit a response in wild type cells. The elevated levels of cytokines but not chemokines the in serum of *Bcl3*^-/-^ mice highlights the importance of MAPK signalling in regulating cytokine synthesis ^4-7^. This relies on MAPK dependent increases in mRNA stability and translation rather than transcription. These findings also suggest that signals that alter the expression of BCL-3, such as cytokines, may also alter the MAPK activation threshold and influence the cellular decision to mount an inflammatory response to TLR stimulation^28^.

In summary, this study identifies the nuclear cytoplasmic shuttling TPL-2 as a novel mechanisms regulating TLR-induced MAPK activity and the MAPK activation threshold. Our data establishes BCL-3 as a unique factor that integrates the regulation of NF-ĸB transcriptional activity and MAPK activation in the nucleus. The possibility of modifying the MAPK activation threshold, through altering TPL-2 localisation, stability or interaction with BCL-3, offers the potential to modify the cellular decision to initiate inflammation for therapeutic benefit.

## Acknowledgments

The authors acknowledge the support of the Medical Research Council (MR/M010694/1) (RC), and the Biotechnology and Biological Sciences Research Council (BB/M003671/1) (RC) and the COST Action BM1404 Mye-EUNITER (www.mye-euniter.eu), supported by COST (European Cooperation in Science and Technology).

## Materials and Methods

### Cell Culture

HEK293T, RAW 264.7 cells and 3T3 mouse embryonic fibroblasts were cultured in DMEM containing 10% fetal bovine serum or bovine calf serum respectively with the addition of, 2mM glutamine, and 100 units/ml penicillin/streptomycin. Bone marrow derived macrophages (BMDM) were prepared *in vitro* as described previously^20^. Briefly, bone marrow was isolated from age and sex matched C57BL6/J mice at 8-12 weeks old, cultured in bacterial petri dish with DMEM containing 10% fetal bovine serum, 2mM glutamine, and 100 units/ml penicillin/streptomycin supplemented with 30% L929 conditioned media for 7 days, with media replacement after four days. BMDMs were re-plated on day 7 into tissue culture treated dishes without L929 conditioned media supplement, rested overnight and experiments were performed on day 8.

### Plasmids, Transfection and reagents

Mammalian expression vectors for BCL-3 and p105 were as previously described^29^. TPL-2-MYC and TPL-2D270A-MYC in pcDNA3 were generous gifts from Stephen Ley ^18^. pcDNA3.1-TPL-2-N-DYK expression vectors were generated by Genscript using TPL-2 cDNA accession number NM_053847. pRevGFP and pRev(1.4)-GFP were generous gifts from B. Henderson and are described elsewhere^27^. The DNA fragment coding for the TPL-2 candidate nuclear export sequence KEEIDLLINHLNVSEVLDIMENLYA was cloned into the pRev (1.4)-GFP vector via BamHI and AgeI sites. The specific oligonucleotides used were as follows : 5’GATCCAAAAGAAGAGATTGA-TTTATTAATTAACCATTTAAACGTGTCGGAAGTCCTGGACATCATGGAGAACCTTTATGC AA3’ and 5’CCGGTTGCATAAAGGTTCTCCAGATGTCCAGGACTTCCGACACGTTTAAATGGTTAATTAATAAATCAATCTCTTCTTTTG 3’ (Integrated DNA Technologies). Transfections were performed using the in vitro transfection reagents: Turbofect Fermentas (293T), Attractene, Qiagen (3T3) and Fugene HD, Promega (RAW264.7) according to manufacturer’s protocols. LPS from *Escherichia coli* 055:B5, LMB from *Streptomyces sp*, MG132 and U0126 mono-ethanolate were purchased from Sigma.

### Western blot and immunoprecipation

Whole cell lysates were extracted from cells suspended in RIPA buffer containing 50mM Tris-HCL pH7.4, 1% NP-40, 0.25% deoxycholate, 150mM NaCl, 1mM EDTA, supplemented with 1mM PMSF, 1mM NaF, 1mM Na3VO4, 2μg/ml aprotinin, 1μg/ml pepstatin and 1μg/ml leupeptin. Nuclear and cytoplasmic extracts were obtained using a Nuclear Extract kit according to manufacturer’s instructions (Active Motif). Lysates were resolved using Tris-Glycine SDS-PAGE, transferred to nitrocellulose membranes and immunoblotted with specific antibodies. For co-immunoprecipation experiments equal amounts of whole cell extracts were precleared for 30 minutes at 4°C with protein G agarose, fast flow beads (Millipore) and immunoprecipitated with primary antibody overnight at 4°C. Agarose pellets were washed three times in RIPA buffer and eluted by incubation at 95°C in 2X sample buffer. For endogenous immunoprecipitation experiments TPL-2 was eluted using the TPL-2 blocking peptide by incubation at room temperature for 10 minutes with occasional agitation (Santa Cruz). Equal volumes of resuspended immunoprecipitates were analysed by western blot. For ubiquitination assays, cells were incubated with 10mM N-ethylmaleamide (N-EM) for 30 seconds and washed in PBS/10mM N-EM. Cells were lysed in 1% SDS, boiled for 5 min and sonicated. Cleared lysates were diluted (1/10) in RIPA buffer supplemented with 20mM NEM. Immunoprecipitation was performed as above and analysed by western blot.

### Antibodies

Anti MYC (sc-40,) anti-TPL-2 M20 (sc-720) and anti-TPL-2 H7 (sc-373677) were purchased from Santacruz. Anti-FLAG rabbit (SAB 4301135), anti-FLAG M2 mouse (F1804), anti-HDAC1 (AV38530), anti α- Tubulin (t6074) and anti-B-actin (SAB 1305567) were purchased from Sigma. Antibodies against p-ERK (#9103), ERK (#9102), p-MEK (#9154), MEK (#4694), pRSK (#23556) and RSK2 (#9340) were purchased from Cell signalling technology and anti Xpress from Invitrogen. Secondary antibodies for Western blotting were purchased from GE healthcare. AF-488 anti-rabbit (A11008), AF-594 anti-mouse (A11005) AF-594 anti-rabbit (A11012) immunofluorescence secondary antibodies were purchased from Life Technologies.

### Luciferase Assay

RAW 264.7 cells were transiently transfected with pAP-1 an AP-1 promoter reporter plasmid (Clontech) and the *Renilla* luciferase expression vector pRL-TK (Promega) for 24 hours. Luciferase activity was measured using the Promega’s dual-Luciferase reporter assay system. For all samples, firefly luciferase activity was divided by that of the *Renilla* luciferase activity to normalize for the transfection efficiency as previously described.

### Gene expression

Total RNA was isolated using RNeasy kits (Qiagen) and reverse transcribed using NanoScript 2 reverse transcription kit (Primer Design). Real-time PCR was performed with SYBR Green SuperMix with ROX (PerfeCTa) using QuantiTect Primer Assays (Qiagen). Data were normalised to TBP and gene expression changes calculated using the 2-∆∆CT method. Targeted RNA-seq analysis was performed using the QIAseq Targeted Mouse Inflammation and Immunity Transcriptome RNA Panel according to manufacturer’s protocols.

### Cytokine analysis

1×10^6^ BMDM were plated per well of a flat bottom 96-well microplate in 200ul of complete media. Cells were left unstimulated or stimulated with 0.1-10ng/ml of LPS for 4 hours. For each experiment supernatants from 3 technical replicates were combined and stored at −80C until analysis. Cytokine concentrations were determined using Bio-plex Pro mouse cytokine Grp1 Panel 23-plex according to manufactures instructions using the Bio-Plex 200 system.

### Immunofluorescence microscopy

WT or *Nfkb1*^-/-^ 3T3s were plated on glass coverslips and transfected for 24 hours. For treatments, cells were incubated with vehicle controls or with 20nM LMB (Sigma) and 20uM MG132 (Sigma) alone or in combination for 2 hours. Cells were fixed and permeablized with ice-cold methanol for 15 minutes, blocked in PBS-0.05% TWEEN (PBS-T) containing 5% BSA. Primary and secondary antibodies were diluted in 1%BSA/PBS-T and staining was performed overnight at 4 C and for 1 hour at room temperature respectively. Nuclear DNA was counterstained using DAPI mounting medium (Vectashield, Vector Laboratories). Immunofluorescent images were acquired using a Zeiss Cell Observer SD confocal fluorescent microscope (Zeiss). BMDM were plated in black 96-well thin bottom (μClear®) microplate. Following LPS stimulations, media was removed and cells immediately fixed with 4% paraformaldehyde, permeabilized with ice-cold methanol and blocked with 5% goat serum in PBS. Primary and secondary antibodies were diluted in PBS-T containing 1% BSA and staining was performed overnight at 4 C and for 1 hour at room temperature respectively, nuclear DNA was counterstained using Hoescht. Images were acquired using confocal fluorescence microscopy (GE Healthcare).

### Kinase assay

For MEK kinase assays, TPL-2 was immunoprecipated from equal amounts of whole cell extracts overnight as described above. Agarose beads were washed 10 times in 1ml of RIPA buffer and washed twice in 1ml MEK kinase assay buffer. Beads were then incubated in 20μl of MEK kinase assay buffer containing 1μg recombinant inactive MEK1 20μl (Millipore) and 2mM adenosine triphosphate (ATP) at 30°C with occasional agitation for 15 minutes. Beads were pelleted at 11,000 *g*, the supernatant was removed, added to 20μl 2X SDS sample buffer and analysed by western blot for phosphorylated MEK. Immunoprecipitates were also eluted from the beads and analysed by western blot.

## Supplementary Data

**Figure S1.**
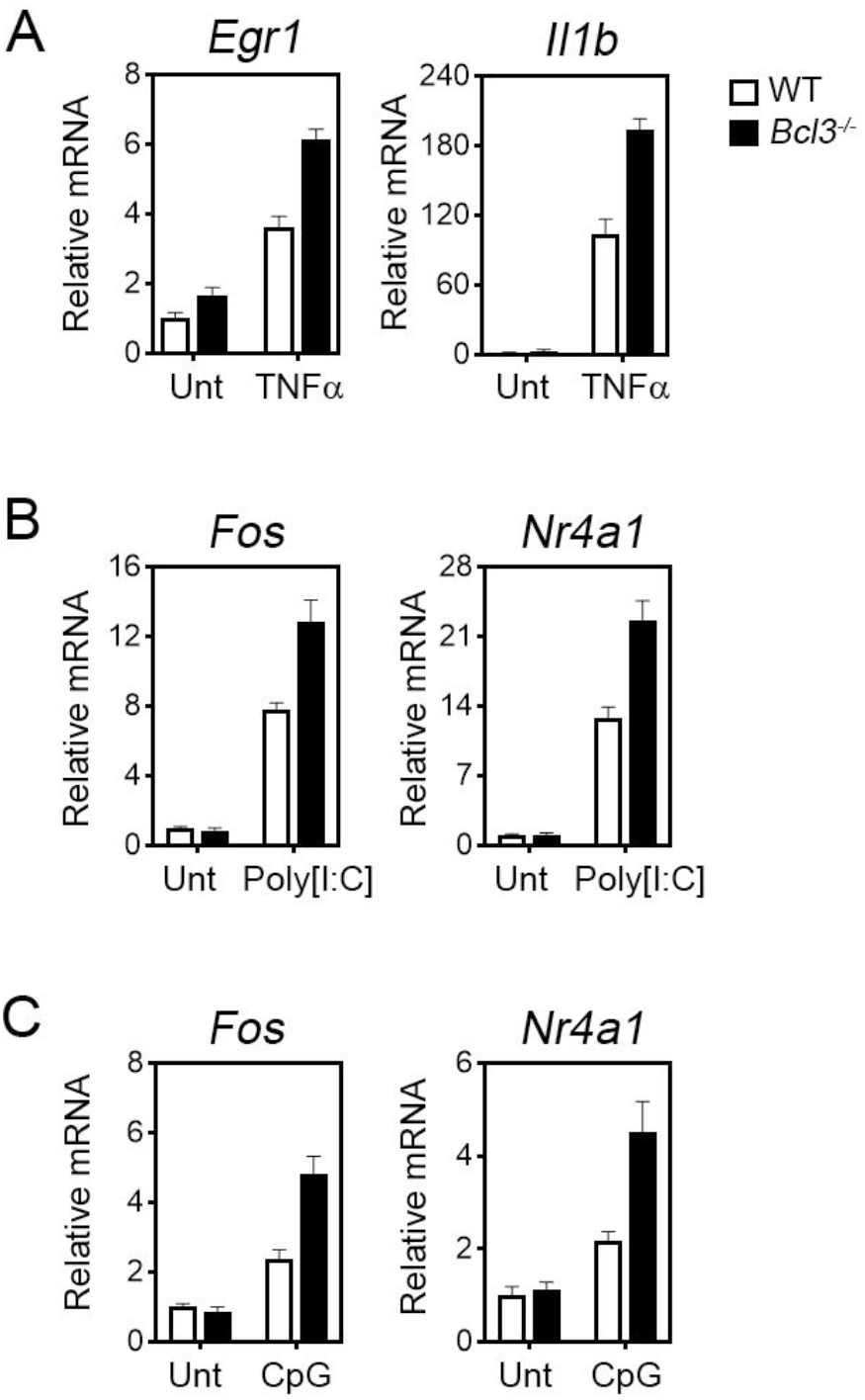
Increased expression of MAPK regulated genes in *Bcl3*^-/-^ macrophages treated with TLR ligands and TNFα. **(A)** WT and *Bcl3*^-/-^ macrophages were stimulated with TNFα (0.2ng/ml for 60 mins) and *Egr1* and *Il1b* mRNA measured by QPCR. **(B)** WT and *Bcl3*^-/-^ macrophages were stimulated with Poly[I:C] (50μg/ml for 60 mins) and *Fos* and *Nr4a1* mRNA measured by QPCR. **(C)** WT and *Bcl3*^-/-^ macrophages were stimulated with CpG (10nM/ml for 60 mins) and *Fos* and *Nr4a1* mRNA measured by QPCR.

**Figure S2.**
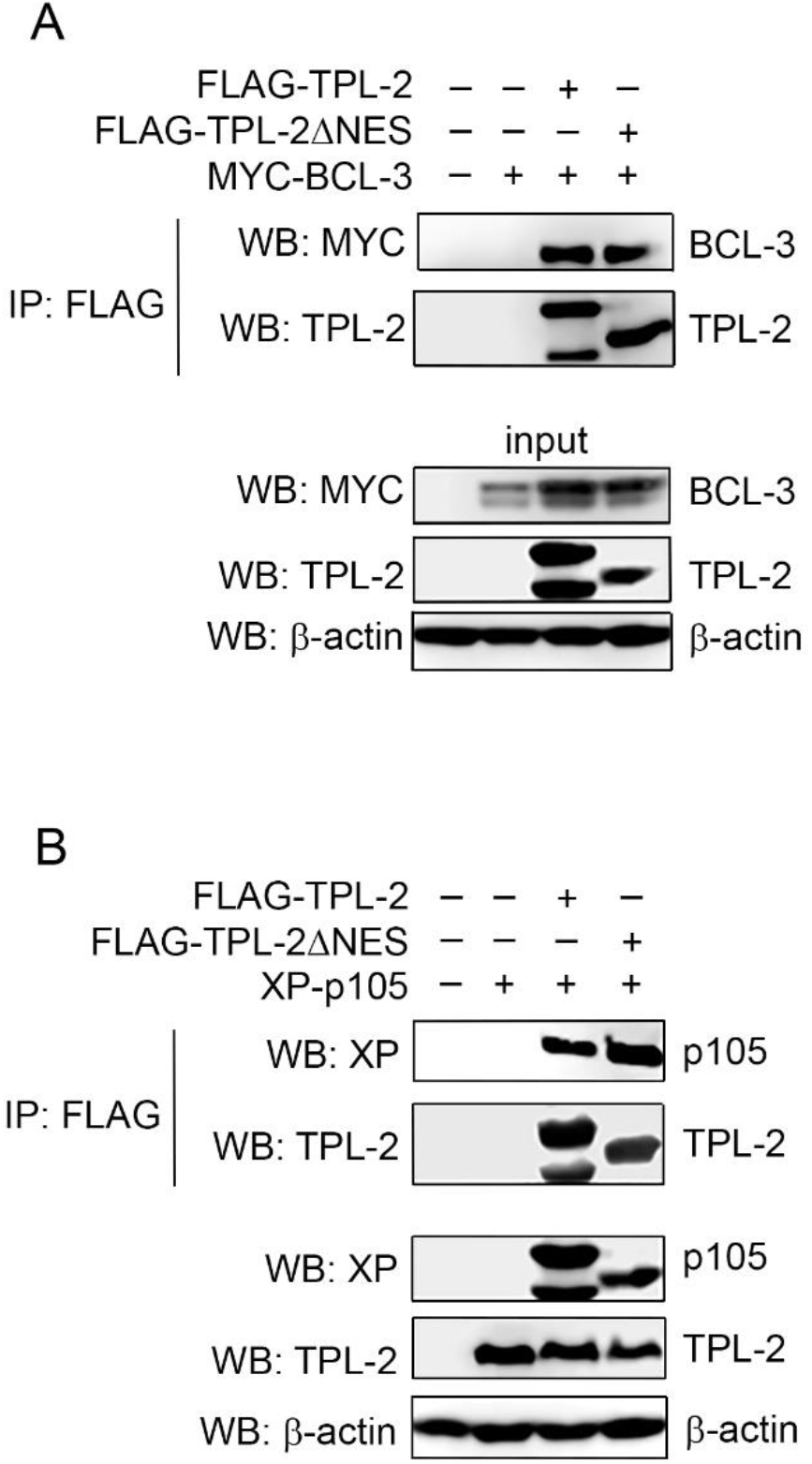
The nuclear export sequence of TPL-2 is not required for interaction with p105 or BCL-3. **(A)** HEK293T cells were transfected with plasmids encoding FLAG-TPL-2, FLAG-TPL-2ΔNES and MYC-BCL-3. TPL-2 and TPL-2ΔNES were immunoprecipitated and interaction with BCL-3 assessed by immunoblot using the antibodies indicated. **(B)** HEK293T cells were transfected with plasmids encoding FLAG-TPL-2, FLAG-TPL-2ΔNES and XP-p105. TPL-2 and TPL-2ΔNES were immunoprecipitated and interaction with p105 assessed by immunoblot using the antibodies indicated.

**Figure S3.**
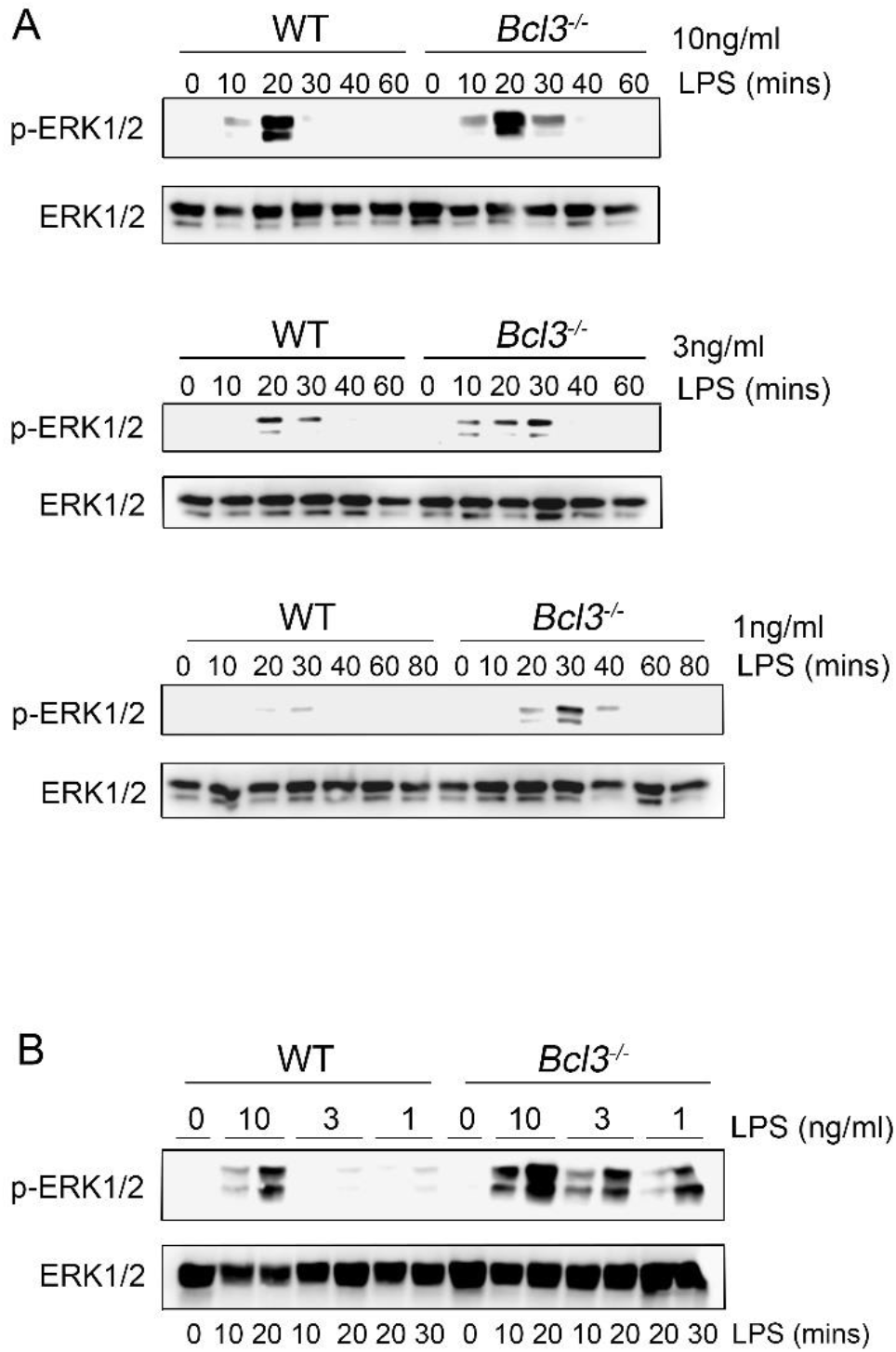
BCL-3 controls the MAPK activation threshold. **(A)** WT and *Bcl3*^-/-^ macrophages were stimulated with the indicated concentrations of LPS for the time points shown and ERK1/2 phosphorylation measured by immunoblot. **(B)** WT and *Bcl3*^-/-^ macrophages treated with the indicated concentrations of LPS for the time points shown and ERK1/2 phosphorylation measured by immunoblot.

**Figure S4.**
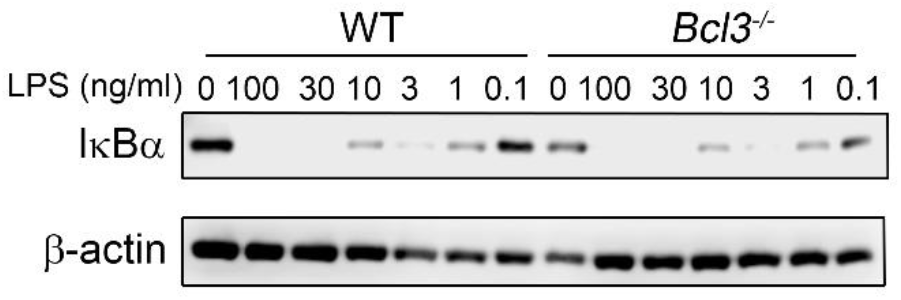
BCL-3 does not alter activation of NF-ĸB. WT and *Bcl3*^-/-^ macrophages were stimulated with the indicated concentrations of LPS and IĸBα levels measured by immunoblot.

**Figure S5.**
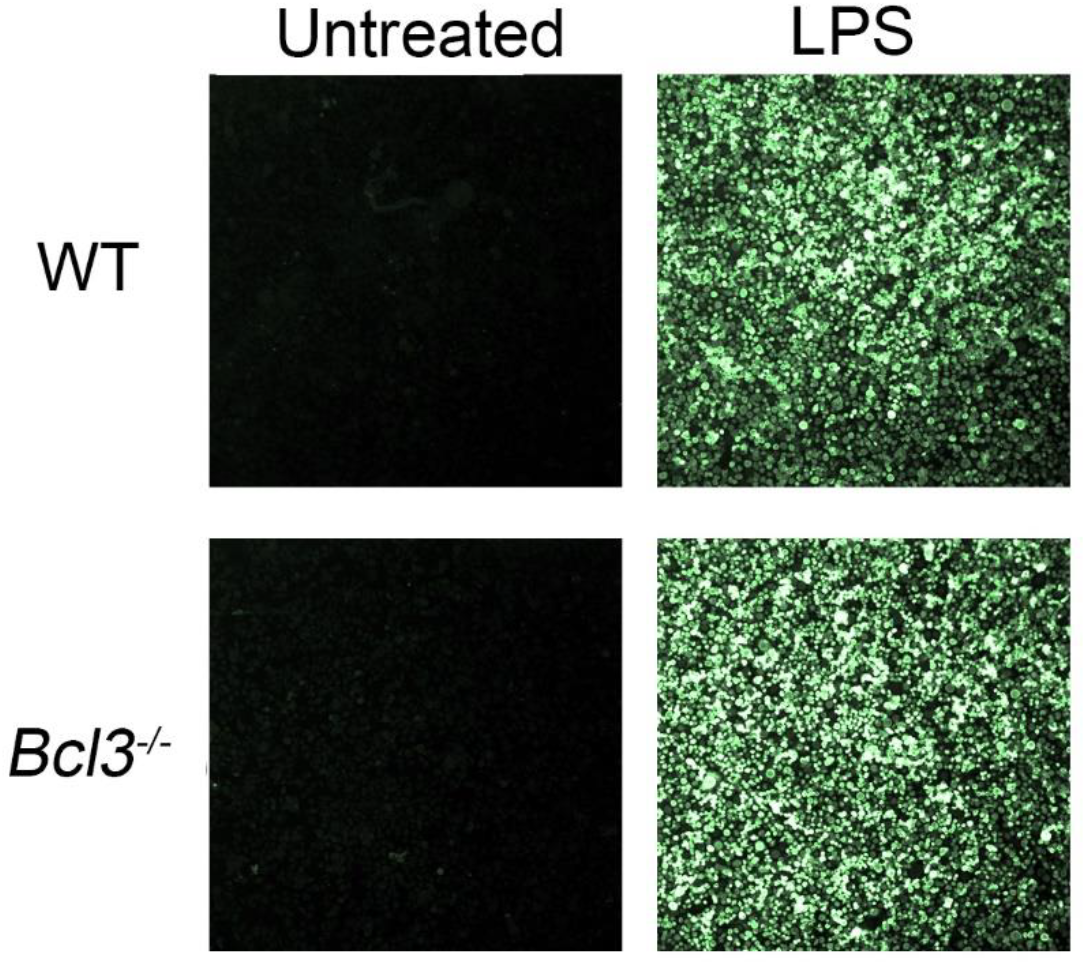
Equivalent numbers of cells WT and *Bcl3*^-/-^ macrophages activate the MAPK pathway in response to LPS. WT and *Bcl3*^-/-^ macrophages were stimulated with LPS (10ng/ml for 30 mins) prior to immunofluorescence staining with anti-phospho-ERK1/2 antibody and imaging by confocal microscopy. Data is representative of three independent experiments.

**Figure S6.**
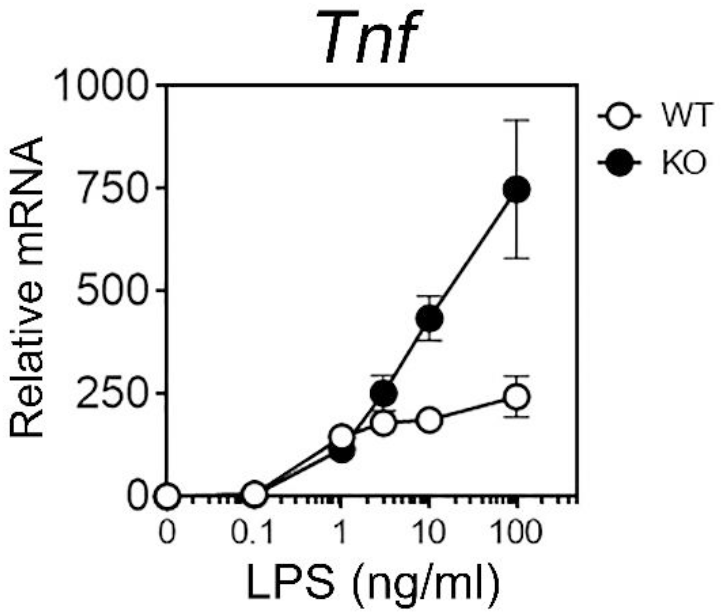
Dose dependent induction of *Tnf* expression in WT and *Bcl3*^-/-^ macrophages. WT and *Bcl3*^-/-^ macrophages were stimulated with the indicated concentrations of LPS for 1 hour prior to analysis by QPCR. Data is representative of 3 independent experiments.

**Figure S7.**
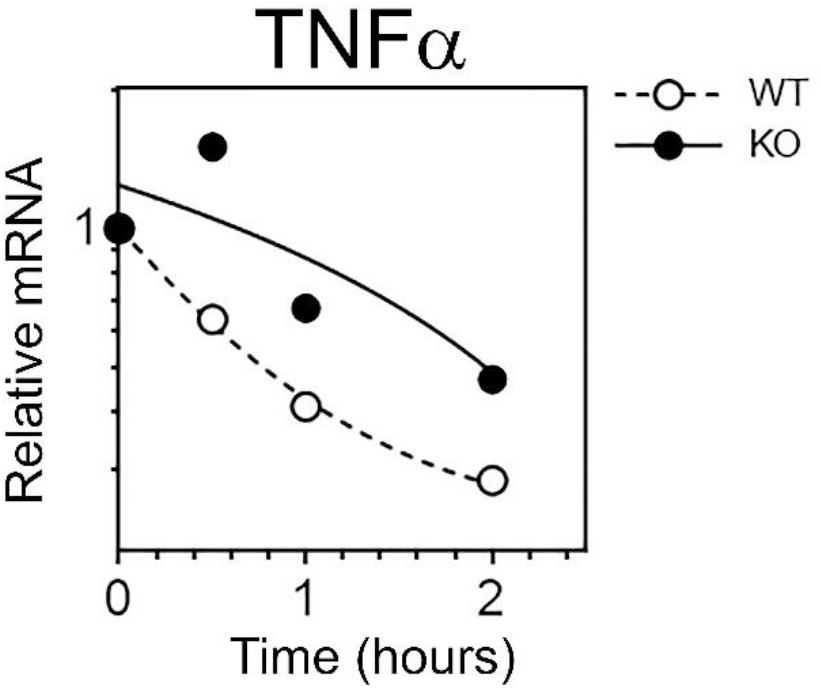
Increased *Tnf* mRNA stability in LPS stimulated *Bcl3*^-/-^ macrophages. WT and *Bcl3*^-/-^ macrophages were stimulated with LPS (10ng/ml) for 30 mins prior to the addition of actinomycin D. *Tnf* mRNA was measured by QPCR at the indicated timepoint following LPS stimulation. Data is representative of three independent experiments.

**Figure S8.**
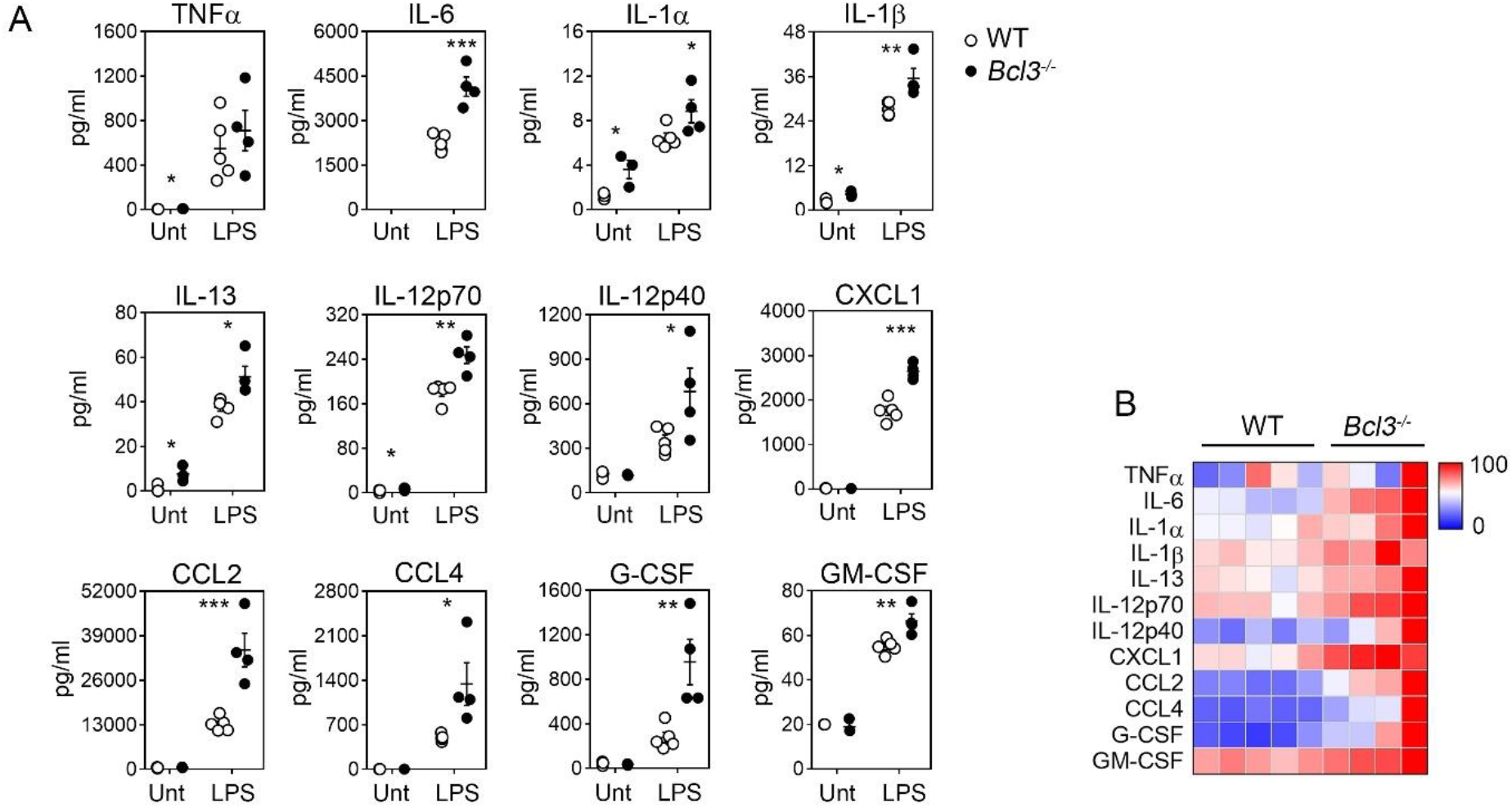
Increased serum cytokine levels in LPS treated *Bcl3*^-/-^ mice. **(A)** Serum from untreated WT (n=3) and *Bcl3*^-/-^ (n=3) mice, and WT (n=5) and *Bcl3*^-/-^ (n=4) mice treated with LPS (2.5 μg/mouse *i.p*) for 1 hour was analysed for the indicated factors. **(B)** Heatmap showing relative levels of serum factors in LPS treated WT and *Bcl3*^-/-^. ****p<0.0001, ***p<0.001, **p<0.003, *p<0.01.

## References

1 Carmody, R. J. & Chen, Y. H. Nuclear factor-kappaB: activation and regulation during toll-like receptor signaling. Cell Mol Immunol 4, 31–41 (2007).

2 Arthur, J. S. & Ley, S. C. Mitogen-activated protein kinases in innate immunity. Nat Rev Immunol 13, 679–692, doi:10.1038/nri3495 (2013).

3 Mitchell, J. P. & Carmody, R. J. NF-kappaB and the Transcriptional Control of Inflammation. Int Rev Cell Mol Biol 335, 41–84, doi:10.1016/bs.ircmb.2017.07.007 (2018).

4 Dumitru, C. D. et al. TNF-alpha induction by LPS is regulated posttranscriptionally via a Tpl2/ERK-dependent pathway. Cell 103, 1071–1083 (2000).

5 Rousseau, S. et al. TPL2-mediated activation of ERK1 and ERK2 regulates the processing of pre-TNF alpha in LPS-stimulated macrophages. J Cell Sci 121, 149–154, doi:10.1242/jcs.018671 (2008).

6 Lopez-Pelaez, M. et al. Cot/tpl2-MKK1/2-Erk1/2 controls mTORC1-mediated mRNA translation in Toll-like receptor-activated macrophages. Mol Biol Cell 23, 2982–2992, doi:10.1091/mbc.E12-02-0135 (2012).

7 Guma, M. et al. Constitutive intestinal NF-kappaB does not trigger destructive inflammation unless accompanied by MAPK activation. J Exp Med 208, 1889–1900, doi:10.1084/jem.20110242 (2011).

8 Karin, M. & Delhase, M. The I kappa B kinase (IKK) and NF-kappa B: key elements of proinflammatory signalling. Semin Immunol 12, 85–98, doi:10.1006/smim.2000.0210 (2000).

9 Lee, R. E., Walker, S. R., Savery, K., Frank, D. A. & Gaudet, S. Fold change of nuclear NF-kappaB determines TNF-induced transcription in single cells. Mol Cell 53, 867–879, doi:10.1016/j.molcel.2014.01.026 (2014).

10 Sung, M. H. et al. Switching of the relative dominance between feedback mechanisms in lipopolysaccharide-induced NF-kappaB signaling. Sci Signal 7, ra6, doi:10.1126/scisignal.2004764 (2014).

11 Ferrell, J. E.Jr., & Ha, S. H. Ultrasensitivity part II: multisite phosphorylation, stoichiometric inhibitors, and positive feedback. Trends Biochem Sci 39, 556–569, doi:10.1016/j.tibs.2014.09.003 (2014).

12 Huang, C. Y. & Ferrell, J. E., Jr., Ultrasensitivity in the mitogen-activated protein kinase cascade. Proc Natl Acad Sci U S A 93, 10078–10083 (1996).

13 Gottschalk, R. A. et al. Distinct NF-kappaB and MAPK Activation Thresholds Uncouple Steady-State Microbe Sensing from Anti-pathogen Inflammatory Responses. Cell Syst 2, 378–390, doi:10.1016/j.cels.2016.04.016 (2016).

14 Gottschalk, R. A. et al. Distinct NF-kappa B and MAPK Activation Thresholds Uncouple Steady-State Microbe Sensing from Anti-pathogen Inflammatory Responses. Cell Syst 2, 378–390, doi:10.1016/j.cels.2016.04.016 (2016).

15 Beinke, S. et al. NF-kappaB1 p105 negatively regulates TPL-2 MEK kinase activity. Mol Cell Biol 23, 4739–4752 (2003).

16 Belich, M. P., Salmeron, A., Johnston, L. H. & Ley, S. C. TPL-2 kinase regulates the proteolysis of the NF-kappaB-inhibitory protein NF-kappaB1 p105. Nature 397, 363–368, doi:10.1038/16946 (1999).

17 Waterfield, M. R., Zhang, M., Norman, L. P. & Sun, S. C. NF-kappaB1/p105 regulates lipopolysaccharide-stimulated MAP kinase signaling by governing the stability and function of the Tpl2 kinase. Mol Cell 11, 685–694 (2003).

18 Beinke, S., Robinson, M. J., Hugunin, M. & Ley, S. C. Lipopolysaccharide activation of the TPL-2/MEK/extracellular signal-regulated kinase mitogen-activated protein kinase cascade is regulated by IkappaB kinase-induced proteolysis of NF-kappaB1 p105. Mol Cell Biol 24, 9658–9667, doi:10.1128/MCB.24.21.9658-9667.2004 (2004).

19 Waterfield, M., Jin, W., Reiley, W., Zhang, M. & Sun, S. C. IkappaB kinase is an essential component of the Tpl2 signaling pathway. Mol Cell Biol 24, 6040–6048, doi:10.1128/MCB.24.13.6040-6048.2004 (2004).

20 Carmody, R. J., Ruan, Q., Palmer, S., Hilliard, B. & Chen, Y. H. Negative regulation of toll-like receptor signaling by NF-kappaB p50 ubiquitination blockade. Science 317, 675–678, doi:10.1126/science.1142953 (2007).

21 Collins, P. E., Kiely, P. A. & Carmody, R. J. Inhibition of transcription by B cell Leukemia 3 (Bcl-3) protein requires interaction with nuclear factor kappaB (NF-kappaB) p50. J Biol Chem 289, 7059–7067, doi:10.1074/jbc.M114.551986 (2014).

22 McNab, F. W. et al. TPL-2-ERK1/2 signaling promotes host resistance against intracellular bacterial infection by negative regulation of type I IFN production. J Immunol 191, 1732–1743, doi:10.4049/jimmunol.1300146 (2013).

23 Yang, H. T. et al. Coordinate regulation of TPL-2 and NF-kappaB signaling in macrophages by NF-kappaB1 p105. Mol Cell Biol 32, 3438–3451, doi:10.1128/MCB.00564-12 (2012).

24 Cho, J. & Tsichlis, P. N. Phosphorylation at Thr-290 regulates Tpl2 binding to NF-kappaB1/p105 and Tpl2 activation and degradation by lipopolysaccharide. Proc Natl Acad Sci U S A 102, 2350–2355, doi:10.1073/pnas.0409856102 (2005).

25 Xu, D. et al. LocNES: a computational tool for locating classical NESs in CRM1 cargo proteins. Bioinformatics 31, 1357–1365, doi:10.1093/bioinformatics/btu826 (2015).

26 Fu, S. C., Imai, K. & Horton, P. Prediction of leucine-rich nuclear export signal containing proteins with NESsential. Nucleic Acids Res 39, e111, doi:10.1093/nar/gkr493 (2011).

27 Henderson, B. R. & Eleftheriou, A. A comparison of the activity, sequence specificity, and CRM1-dependence of different nuclear export signals. Exp Cell Res 256, 213–224, doi:10.1006/excr.2000.4825 (2000).

28 Palmer, S. & Chen, Y. H. Bcl-3, a multifaceted modulator of NF-kappaB-mediated gene transcription. Immunol Res 42, 210–218, doi:10.1007/s12026-008-8075-4 (2008).

29 Collins, P. E. et al. Mapping the Interaction of B Cell Leukemia 3 (BCL-3) and Nuclear Factor kappaB (NF-kappaB) p50 Identifies a BCL-3-mimetic Anti-inflammatory Peptide. J Biol Chem 290, 15687–15696, doi:10.1074/jbc.M115.643700 (2015).

